# Standard-GEM supports reproducible development of genome-scale metabolic models

**DOI:** 10.1101/2023.03.21.512712

**Authors:** Mihail Anton, Eivind Almaas, Famke Bäuerle, Moritz E. Beber, Rui Benfeitas, Sara Benito-Vaquerizo, Lars M. Blank, Frederick Clasen, Claudia de Buck, Andreas Dräger, John M. Hancock, Vassily Hatzimanikatis, Matthias König, Dimitra Lappa, Feiran Li, Ulf W. Liebal, Hongzhong Lu, Hongwu Ma, Radhakrishnan Mahadevan, Costas D. Maranas, Adil Mardinoglu, Jens Nielsen, Juan Nogales, Marco Pagni, Jason A. Papin, Jonathan L. Robinson, Benjamín J. Sánchez, Saeed Shoaie, Vetle Simensen, Maria Suarez-Diez, Snorre Sulheim, L. Thomas Svensson, Bas Teusink, Wanwipa Vongsangnak, Ahmad A. Zeidan, Eduard J. Kerkhoven

## Abstract

The field of metabolic modelling at the genome scale continues to expand, with increasing numbers of models being reconstructed, shared and curated. This expansion creates a need to adopt common principles regarding transparency and versioning, in addition to standardisation of file formats, annotations, and model testing. Here, we present a repository standard and template for Git-based genome-scale metabolic models (GEMs) supporting both newly reconstructed and actively curated models, following FAIR principles (findability, accessibility, interoperability, and reusability), and incorporating research software best practices. `Standard-GEM` exposes GEMs across web services and platforms in the metabolic modelling field and enables automatic validation of GEMs. The use of this template for new models and its adoption reduces the effort required to share, validate and reuse GEMs.

## Introduction

Genome-scale metabolic models (GEMs) have been used to understand and guide modifications of metabolism with a wide range of applications, from elucidating disease mechanisms and identifying biomarkers to optimising strains for producing valuable chemicals (Chen et al., 2019, Nielsen and Petranovic, 2025). However, the size of GEMs and the iterative curation work required to build and refine them, as detailed in a 96-step standard operating procedure by Thiele and Palsson (2010), naturally lead modellers to desire a framework inspired by research software best practices. GEM reconstruction and maintenance require extensive iterative curation, including the integration of literature evidence, genome annotation, biochemical knowledge, simulation constraints, and model-quality checks. Consequently, many GEMs are not static research outputs but evolving computational resources that continue to change after initial publication.

Scientific results, including those from model analysis, are increasingly evaluated for reproducibility, the benefits of which have been previously established (Baker, 2016, Blinov et al., 2021, Malik-Sheriff et al., 2020). Reproducibility can be viewed as part of a wider need for open science data management aimed at the reuse of digital research objects (DROs). The FAIR Principles (findability, accessibility, interoperability, reusability) provide an easily accessible suite of recommendations for the sharing of DROs (Wilkinson et al., 2016). These principles are conceived as broadly applicable not only to data but also to other classes of DROs including software tools, workflows, and systems biology models. To operationalise these principles for systems biology models, Research Data Alliance FAIR indicators have been adapted specifically for COMBINE models and archives (Balaur et al., 2025, Bergmann et al., 2014). More recently, the CURE principles (credibility, understandability, reproducibility and extensibility) have been proposed to complement FAIR specifically for computational models of biological systems, addressing aspects of model quality that FAIR alone does not capture (Sauro et al., 2026). As highlighted by the Infrastructure for Systems Biology Europe (ISBE), “sharing [data and] models solely through supplementary material is still common practice” (Stanford et al., 2015), with the recommendation that platforms should be used to disseminate assets. In line with this, EMBO Reports and other scientific journals have advocated using BioModels (Malik-Sheriff et al., 2020) as a default deposition database for sharing published models in standardised formats, such as the Systems Biology Markup Language (SBML) (Keating et al., 2020). ISBE in cooperation with ERASysAPP initiated a key step towards promoting FAIRification in systems biology with the development of FAIRDOM and FAIRDOMHub (Wolstencroft et al., 2017). FAIRDOMHub functions as a publishing platform for systems modelling projects which links their various elements from data to model. These may be located at different physical locations and in different types of repositories, such as SEEK (Wolstencroft et al., 2015) and openBIS (Barillari et al., 2016). Here, we propose to continue this advancement in FAIRification through the implementation of open-source standards both during model creation and in its subsequent curation, not only for making the final model available in a reproducible format, but also enabling reproducibility throughout the model development and curation process (Shin et al., 2023, Tiwari et al., 2021). Deposition of a final model file does not always capture the full development context of a GEM, including the code, data, intermediate decisions, validation history, issues, and pull requests that shaped the model.

This limitation is particularly relevant for community-developed models and models that are updated over multiple releases. The importance of versioning has been well-recognised for biological models and repositories, especially for models that constantly evolve (Beard et al., 2009, Domenzain et al., 2021, Miller et al., 2011). Code versioning can improve the reproducibility and transparency of the curation process (Aite et al., 2018), which begins during the original GEM reconstruction process and ideally continues during the model lifetime whenever new knowledge becomes available, analogous to continuous development in software engineering (Ravikrishnan and Raman, 2015). For example, the consensus GEM of *Saccharomyces cerevisiae* has been developed since 2008 (Herrgård et al., 2008) and continually updated with new curations from different research groups, reaching version 9 in 2024 (Zhang et al., 2024). Evaluation metrics for annotation of model components have been consistently improved across versions (Domenzain et al., 2021). At the same time, models hosted by databases such as the BiGG database (Norsigian et al., 2020) have been updated by applying the curation tool ModelPolisher (King et al., 2016). However, the changes that have been applied to the BiGG models are not traceable by the research community. Such missing provenance information makes it difficult to evaluate changes between versions. Similarly, the latest version of BioModels “transparently tracks changes to the model and associated files behind the scenes” (Malik-Sheriff et al., 2020), but such changes are not readily available on the website as of August 2026. To overcome these challenges, we aim to structure the post-publication manual curation process by making all changes equally explicit and trackable, previously identified as desirable characteristics by Waltemath et al. (2013). Several GEM resources already use Git-based workflows, but the field lacks a minimal repository standard that defines how model files, code, data, documentation, releases, and validation outputs should be organised for reproducible curation.

## Results

Here, we present `standard-GEM` as a lightweight repository standard for reproducible GEM development, together with a template repository and an automated validation pipeline that operationalise the standard (Fig 1). To address the above-mentioned issues of reproducibility and availability of provenance information and apply FAIR principles to GEMs, we establish a specification for model reuse while relying on repository-hosting solutions. Specifically, our approach leverages a fixed folder structure and the use of a Git-based workflow for version control, enhanced by public issue tracking and discussion forums to further boost the documentation process. To this end, the standard introduced by standard-GEM, defined through the .*standard-GEM*.*md* file, is explicit and transparent. Upon this standard, further validation is achieved through an open-source automated pipeline.

**Fig. 1.**
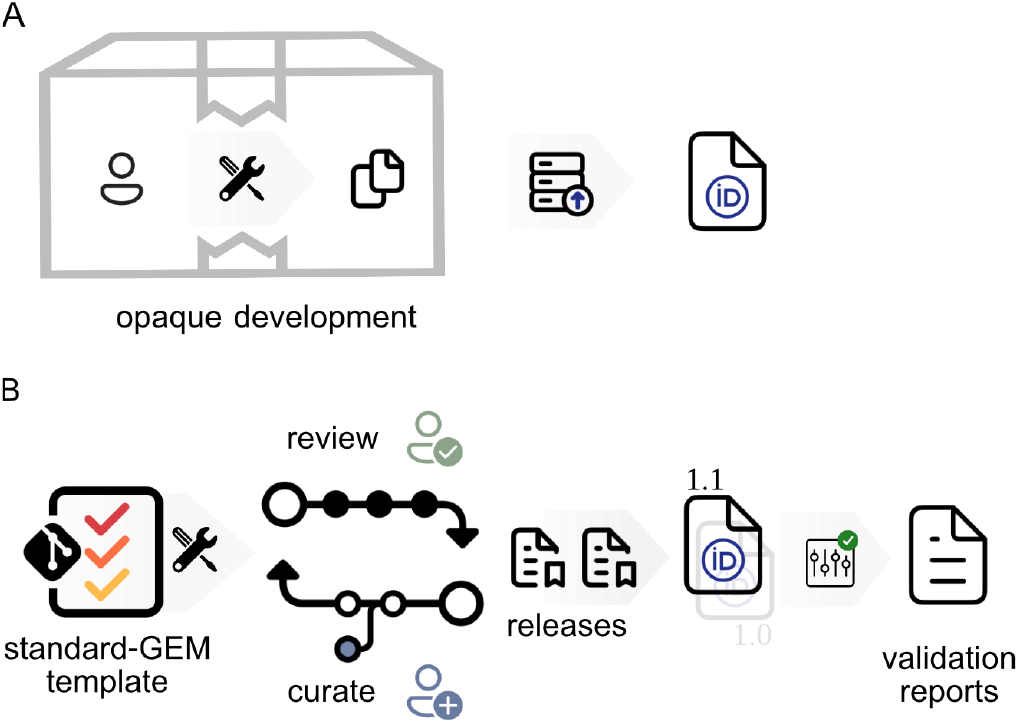
Repository-centred GEM development with standard-GEM. (A)In a conventional deposition-centred workflow, the final model file may be shared through supplementary material, a model database, or an archived release, but the code, data, intermediate versions, validation outputs, and curation rationale are often unavailable or disconnected from the model. (B)Standard-GEM defines a repository-centred workflow in which model files, code, data, documentation, release metadata, and curation history are preserved in a version-controlled repository. The repository can be validated automatically, archived with persistent identifiers, and reused by model databases or web services.

### Template repository

As part of the repository template available at github.com/MetabolicAtlas/standard-GEM, the key file .*standard-GEM*.*md* (Supplementary Note 3) contains all the expectations of the standard, including steps for repository creation, Git workflow, and model file formats. Additionally, using the standard-GEM template repository initiates a well-defined folder structure, including a structured *README* file that facilitates the presentation of the model, and a default licence file (CC BY 4.0).

As a template repository, there are no restrictions on making repositories public. This template can be used to create both private and public repositories identically in GitHub, or in any Git-hosting service. As some functionality of the template uses the GitHub configuration via the .*github* folder, adopting it with another hosting service would entail additional work.

### Automatic validation

GEMs must be maintained to maximise their value. This continuous effort is essential, and it also applies to GEMs created from the standard-GEM template. One of the costs associated with maintaining an open-source GEM is the burden of running checks, ranging from basic sanity checks to more complex evaluations of model content. To reduce this burden, we have created an automatic validation pipeline, publicly available at github.com/MetabolicAtlas/standard-GEM-validation. The pipeline runs on a scheduled basis, which means validation results are updated as the models continue to evolve. This reduces the need for modellers to maintain identical validation pipelines separately, thus reducing the maintenance cost, while keeping the benefit of validation checks. All the results of the pipeline are versioned in JSON format in the same repository as the pipeline and are made available for querying through the website metabolicatlas.github.io/standard-GEM-validation in order to be easily reused by other services. An example of an up-to-date rendering of this JSON is available through the Standard GEMs page at Metabolic Atlas metabolicatlas.org/gems/standard-gems (Supplementary Figure S1).

### Adoption by GEM repositories

The `standard-GEM` topic and the associated validation pipeline also provide a lightweight mechanism to monitor uptake of the standard across the community. At the time of analysis, the pipeline identified 26 GEM repositories adopting standard-GEM, spanning microbial, fungal, animal, and human metabolic models, including both newly reconstructed models and established community models such as yeast-GEM and Human-GEM (Table 1). The associated publications include institutions in multiple countries, illustrating that repositories adopting standard-GEM are linked to geographically distributed modelling efforts (Fig 2).

**Table 1.**
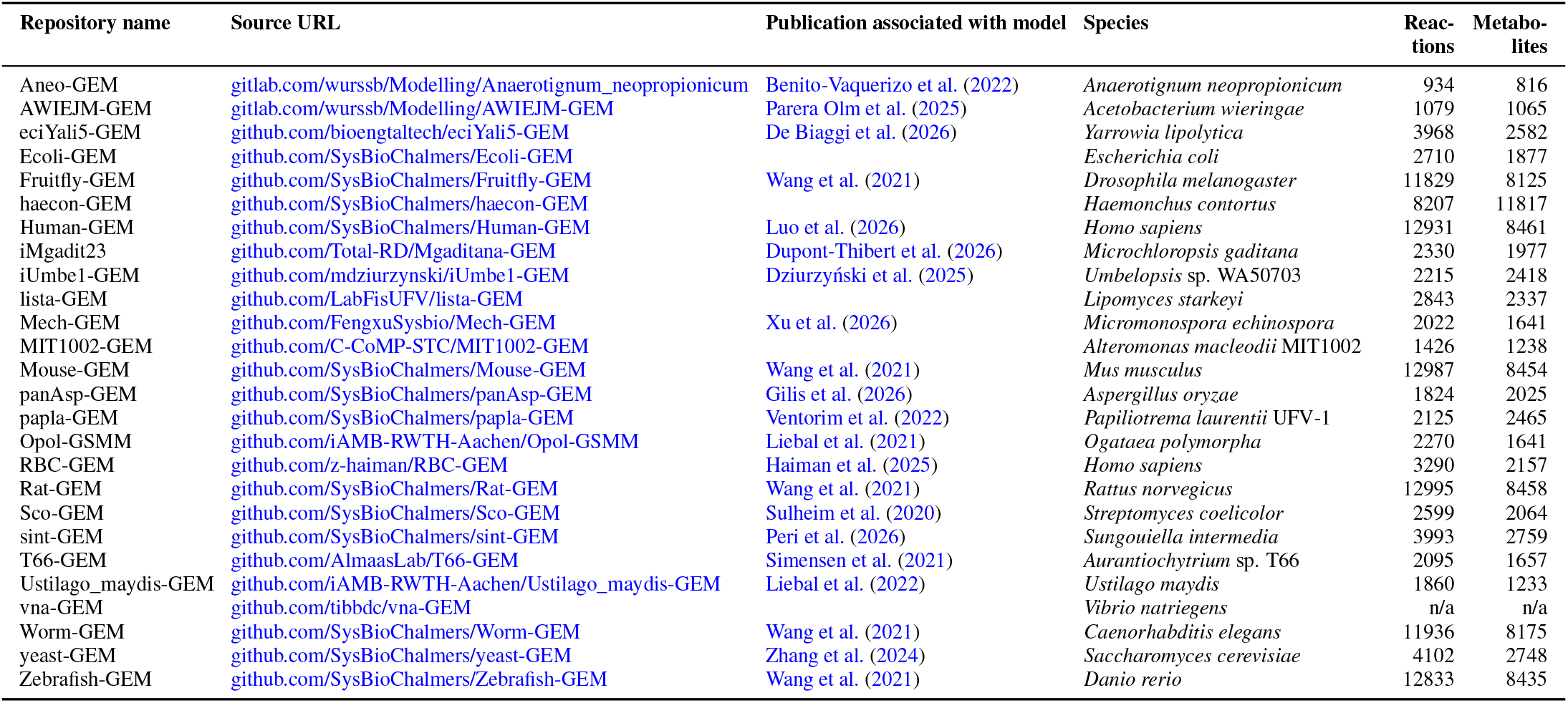
Overview of standard-GEM adoption. Repositories were included if they contained ‘.standard-GEM.md’ and either carried the ‘standard-gem’ topic or were listed in the standard-GEM validation source list. Repository metadata and validation status were obtained from metabolicatlas.github.io/standard-GEM-validation. Publication links were assigned manually from the repository citation metadata or associated publication. Repositories with blank publication fields had no peer-reviewed publication identified at the time of assessment on 1 October 2026.

**Fig. 2.**
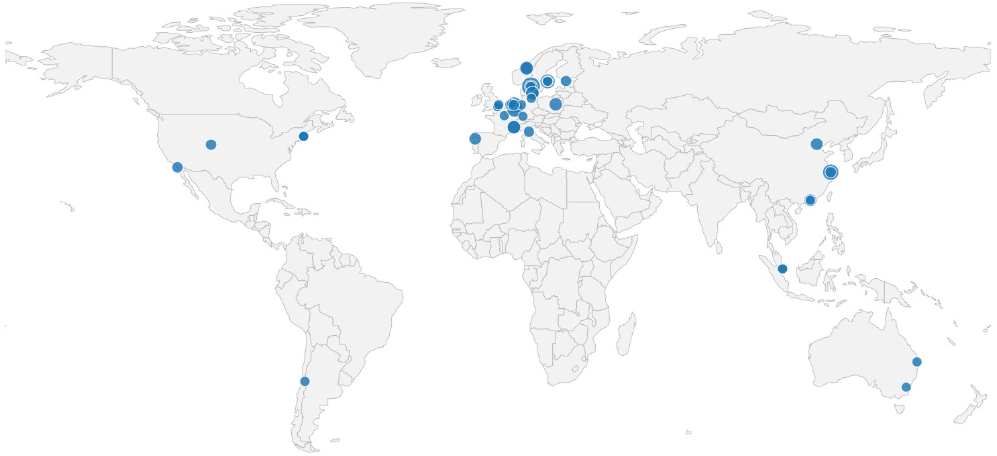
Geographic distribution of affiliations from publications associated with standard-GEM repositories. Dots indicate institutional locations extracted from author affiliations in peer-reviewed publications associated with the GEM repositories listed in Table 1. Larger dots indicate multiple publication authors at the same location. The map represents the geographic distribution of publications linked to repositories adopting standard-GEM, not necessarily the current locations of repository maintainers or all contributors to those repositories.

## Discussion

### Generic infrastructure versus specialised databases

We advocate using standard-GEM through an independent platform that comes at no cost to the scientific community. While a Git-based standard is a platform-independent solution, closer integration with GitHub specifically brings increased simplicity, for example, by instantiating a standard-compliant configuration with a single mouse click; maintainers must complete the repository-specific requirements. Nonetheless, as standard-GEM relies on a single plain-text file, it can be applied to any other platforms of distributed version control and source code management such as GitLab or Bitbucket.

A fundamental debate that underpins standard-GEM is the use of a no-cost generic infrastructure versus a specialised and non-free platform. MEMOsys (Pabinger et al., 2011) provides an instructive precedent for model-versioning infrastructure. It demonstrated the value of centralised version control for metabolic models and provided access to model development history. Its database-centred design, however, differs from the repository-centred approach taken here. In MEMOsys, model properties are mapped into a dedicated database infrastructure; in standard-GEM, the model files, code, data, documentation, issues, pull requests, releases, and validation outputs remain co-located in a general-purpose version-controlled repository. The two approaches are therefore complementary rather than mutually exclusive: database-centred systems can provide curated access and query functionality, whereas repository-centred systems preserve the development context and curation provenance close to the primary research artefacts.

Instead of storing all properties of a metabolic model in a database, standard-GEM provides a standardised format such that researchers can continue to work in an open-source way, directly combining model with code, while keeping the output file format easy to reuse by different model-centric websites such as BioModels, BiGG, FAIRDOMHub, Metabolic Atlas (Wang et al., 2021), and JWS Online (Olivier and Snoep, 2004). ModelBricks defines an alternative infrastructure of small models that requires the development of a database, tools, and content (Cowan et al., 2019). To some extent, standard-GEM facilitates the reuse of a model by forking the source repository. However, how to combine multiple `standard-GEMs` as proposed through ModelBricks remains an open question.

Compared to other standardisation efforts and initiatives, standard-GEM brings many, if not all, advantages without requiring the community to maintain a dedicated central model-versioning infrastructure. While the reliance on Git might have been considered a hindrance, Git has become ubiquitous in bioinformatics training programmes, thus making standard-GEM a practical tool for the target user, reducing the need for additional training.

### Suitability of a Git-based workflow

The use of Git in computational biology has become increasingly popular, especially via hosting platforms like GitHub and GitLab (Cadwallader et al., 2022, Perkel, 2016), including among genome-scale metabolic modellers (Supplementary Table S1). Introductions to how to use Git and GitHub have been previously published (Blischak et al., 2016, Perez-Riverol et al., 2016).

A good standard for GEM versioning should facilitate answering the 7W questions of provenance (*What, Who, When, Where, Which, Why, With/How*) that underpin each action in the curation process (Ram and Liu, 2012). One approach to address provenance post-curation is with algorithms for difference detection in models of biological systems (Hari et al., 2024, Scharm et al., 2016, Scott-Brown and Papachristodoulou, 2017). However, these still leave “scope for further extensions to provide hypotheses for the Why” (Scharm et al., 2016).

When used with disciplined commit messages, pull requests, and issue templates, a Git-based workflow can support the 7W provenance framework. Hosting a Git-based project on an online platform such as GitHub makes these answers transparent by showing *What* the change is (a Git commit), *Who* made the change (recorded author or committer), *When* it occurred (Git commit timestamp), *Where* it occurred (in which file or part of the model), *Which* changes were made (visible through a Git diff). The *Why* and *With* are addressable through the functionality of online platforms, e.g., commit messages and GitHub’s issues and pull requests for which standard-GEM provides templates. Discussions also occur openly on the GitHub platform, with changes to the standard being connected to issues, which can be raised by GitHub users. Therefore, if used adequately, Git-based versioning for models can bring benefits in terms of traceability. Through this approach, standard-GEM aims to document all changes of the standard and to equally invite contributors.

Another advantage of using Git-versioned GEMs is that it makes comparisons between consecutive versions of the model straightforward. With Git-flow, a common development workflow on Git-based platforms, the differences across versions can be easily inspected via branch comparisons.

However, one limitation inherent to the use of Git is that line order changes matter, even if the lines’ content is otherwise identical. The recommendation is that tools maintain the order of model components to provide a consistent order and robustness to the model files and make them suitable for Git. Maintaining the order can be achieved if tools apply the same sorting to the model components, even if components are added or removed.

While promising, the use of code versioning principles when working with GEMs is not entirely straightforward. One can look again at research software, where best practices have been defined and widely spread during the past decade. At the same time, certain code conventions and configuration options can become overly complex for metabolic modelling projects. MEMOTE addresses model testing and also provides `memote-cookiecutter`, a utility for instantiating a Git-versioned project structure (Lieven et al., 2020). This has been important for promoting systematic testing and standardisation of GEM projects. Here, standard-GEM differs in emphasis: it defines a minimal, explicit repository specification that can be inspected directly in `.standard-GEM.md` and adopted without committing to a restrictive project scaffold. The aim is therefore not to replace MEMOTE or `memote-cookiecutter`, but to provide a lightweight repository convention that can incorporate MEMOTE-based testing where appropriate.

Overall, standard-GEM leverages many advantages of Git-based version control, while keeping the approach transparent and lightweight.

### Adhering to standards

At its core, standard-GEM is a standard as per the definition that it is based on explicit community agreement, and that “introducing standards should not be a goal in itself, but should help [… ] solve problems” (Brazma et al., 2006). This minimalist approach is similar to the contributing guidelines that GitHub encourages via `CONTRIBUTING.md`.

A key principle has been to build on existing standards and reduce complexity. As noted by Yurkovich et al. (2017), “one of the biggest hurdles for those attempting to link different pieces of software is standardisation”, to which they formulate “Lesson 6: Adopt or Develop Standards.” In line with this recommendation, standard-GEM limits the development of new standards to where they are strictly necessary, e.g., by promoting model file formats in the community. A prominent example is SBML as the *de facto* file format for GEMs. Other file formats exist, such as SBtab (Lubitz et al., 2016), which provides a spreadsheet-like interface to the model and its annotation. Different modelling formats have been reviewed by Dräger and Palsson (2014) and Schreiber et al. (2020), and usage of file formats has been polled by Carey et al. (2020), which informed the file format recommendations in standard-GEM. Furthermore, the repository convention is independent of programming language; compatibility with individual tools depends on the model formats and features they support. A set of features that defines a “gold standard” for metabolic network reconstruction has been compiled regarding content, annotation, and simulation capabilities (Carey et al., 2020). The proposed standards for the *de novo* reconstruction phase are implemented through a templated *README* file that is part of the default standard-GEM configuration. For the proposed curation process, standard-GEM leverages Git-flow and features of Git-based platforms — commit messages, branches, issues, pull requests — to achieve the desired transparency in the reconstruction and curation process (Heavner and Price, 2015). Regarding the proposed standards for simulations, standard-GEM encourages the separation of concerns by using a fixed folder structure and *README* files. Moreover, standard-GEM aims to work towards full compatibility with the COMBINE archive standard (Bergmann et al., 2014) and RO-Crate (Soiland-Reyes et al., 2022). Overall, standard-GEM functions not as a competing standard, but as a pragmatic integration layer that operationalises existing community standards in routine GEM development and curation.

### FAIR

With its reliance on mature technologies and platforms, standard-GEM provides a FAIR-by-design approach to model building, development and curation that is new to the field (Fig 3), while at the same time supporting key principles beyond FAIR.

**Fig. 3.**
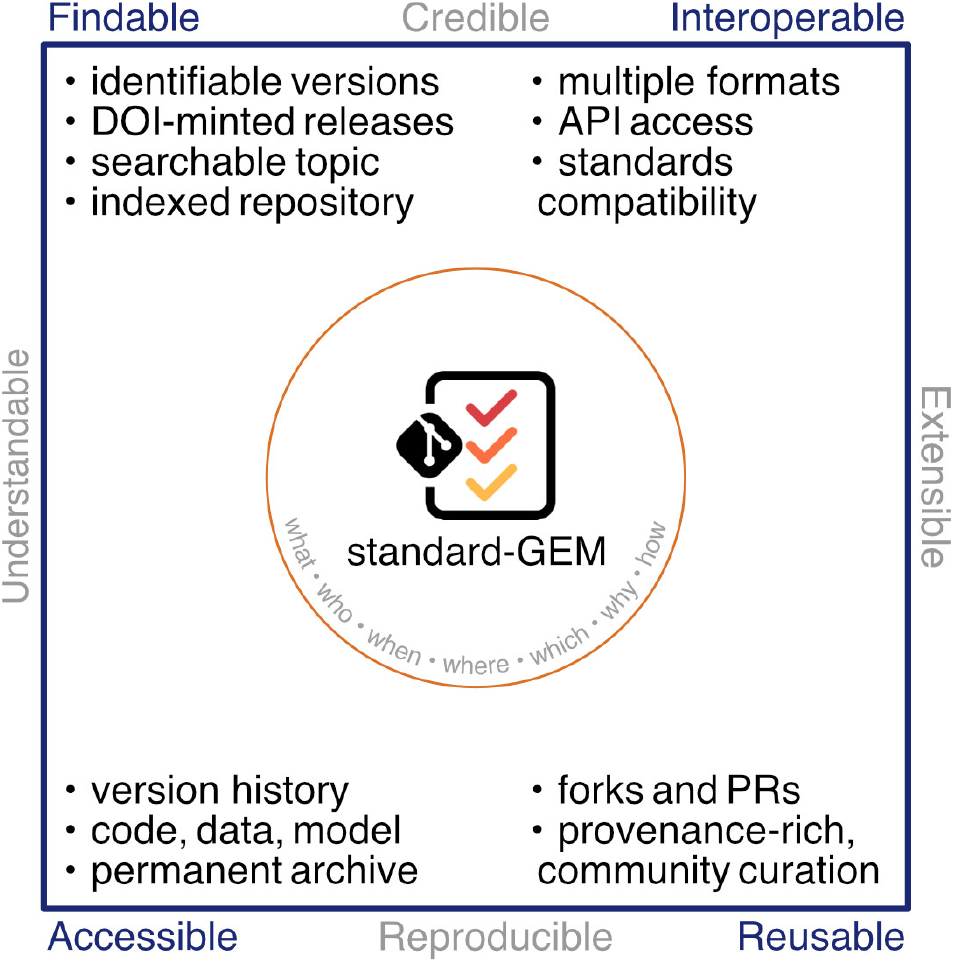
Relationship between standard-GEM, FAIR, and CURE principles. Standard-GEM primarily operationalises FAIR principles at the repository level by improving the findability, accessibility, interoperability, and reusability of GEM research objects. The same repository-level features also support CURE-aligned model practice: validation outputs and transparent curation support credibility; standardised documentation supports understandability; versioned code, data, and model files support reproducibility; and forks, issues, and pull requests support extensibility.

### Findable

Using standard-GEM promotes findability. As mentioned previously, several established resources already exist for storing models that are considered finished, i.e., ready for publication. However, model-file deposition alone may leave accompanying development resources and the rationale for modelling decisions difficult to find. A standard-GEM repository organises these resources alongside the model files, increasing the findability of the information that accompanies an individual model file as a product.

Another aspect of findability to be considered is that of the repositories themselves. To emulate repository categories, GitHub uses searchable topics, and the newly introduced “standard-GEM” topic makes it easy to find all such GEMs on GitHub (Table 1). Additionally, the recommended use of Zenodo ensures automatic DOI minting for each release in the GitHub repository (Supplementary Note 3).

The built-in versioning features make it straightforward to uniquely identify a timestamp-based state of the repository in its entirety – not only the model, but all the files and all the history to that point – through a commit hash, tag, or via the web browser through a release. Moreover, Git hosting platforms provide ways of finding and inspecting repository forks, such as when another group builds upon the work of the original model creators.

### Accessible

The main contribution of standard-GEM to FAIRification is via accessibility. When depositing a model file in a deposition database, only a minor part that goes into the development of the model becomes accessible at that resource. For example, model-only deposition routes do not necessarily capture all code, data, and provenance used during model development. Moreover, thanks to the Git versioning system and the online hosting of the repository, even previous versions of the files in the repository remain accessible. Furthermore, in anticipation of the low risk that the repository becomes inaccessible, as part of the .*standard-GEM*.*md* file, we recommend the permanent archiving service Zenodo which integrates seamlessly with GitHub (Supplementary Note 3). In addition, GitHub allows easy switching between the private/public state of a repository, independently of it being instantiated from a repository template such as standard-GEM.

### Interoperable

While SBML has become the *de facto* file format for GEMs, different use-cases where SBML is less optimal have been identified. For example, due to intrinsic properties of the XML format that SBML is based on, even small model curations can lead to more line changes than in other file formats. Instead of promoting a single solution as is usually the case with deposition databases, standard-GEM coordinates the use of established community standards. This is achieved by recommending different file formats, thus increasing the interoperability of the models, which is a key pillar of standard-GEM. The RAVEN toolbox, for example, provides the wrapper function *exportForGit* that exports multiple formats at once, to reduce the chance of out-of-sync exports.

### Reusable

Standard-GEM adds a provenance-centred dimension to reusability. Whereas the use of standard file formats primarily covers reusability per its official interpretation, especially regarding annotation, the provenance sub-principle (FAIR R1.2) is largely overlooked. With a code versioning system, however, changes in a repository tend to appear gradually. In a standard-GEM, incremental curations, e.g., Human-GEM issue 1009 available at https://github.com/SysBioChalmers/human-GEM/issues/1009, have even more context than generic curation databases, such as APICURON (Hatos et al., 2021), which focuses more on curation as a trackable activity than on the curation rationale. Moreover, provenance records for individual curations can support transfer of those curations to related GEMs or databases.

A different aspect of reusability inherent to a standard-GEM is that it is expected to contain code. Large curations, even if implemented manually, benefit from being code-aided. Even if such code is meant for a one-time use, versioning it in the repository makes it reusable for future curations.

Lastly, the reusability of a repository increases once it becomes public on a Git-hosting service. Public repositories can be forked by other users, enabling them to duplicate the entire repository under their account, including the change history. Later, any new changes can be conveniently incorporated in the original repository via a pull request, thus making even work outside the original repository reusable.

### Beyond FAIR

Standard-GEM also supports several aspects of the CURE principles for computational models. It contributes to credibility by encouraging explicit documentation, public issue tracking, validation outputs, and versioned releases. It supports understandability by prescribing a standard repository layout, README structure, and human-readable model representations. It improves reproducibility by preserving model files, code, data, release history, and validation outputs in one version-controlled research object. Finally, it supports extensibility by allowing repositories to be forked, modified, validated, and reintegrated through pull requests. While standard-GEM does not by itself guarantee biological correctness or numerical reproducibility, it does provide the repository-level structure needed to make such assessments inspectable.

### Governance and versioning of the standard

We distinguish between the version of the standard, the version of a GEM repository adopting the standard, and the version of the validation pipeline. By design, standard-GEM is intended to evolve as community practice changes, but such evolution must remain controlled to preserve compatibility across repositories. Changes to the specification are therefore managed through the public standard-GEM repository, where proposed modifications can be discussed as issues or pull requests. Minor changes include clarifications, documentation improvements, and additional non-breaking recommendations. Major changes include new mandatory requirements, changes to required file locations, or altered validation logic that may affect whether existing repositories are considered compliant. Such changes should be introducedthrough versioned releases of the standard and accompanied by migration guidance for existing repositories.

The current version of standard-GEM should therefore be interpreted as a stable implementation of a minimal specification rather than a closed final standard. This governance model allows standard-GEM to remain lightweight while providing a clear mechanism for community input, backwards-compatible improvements, and transparent handling of breaking changes.

### Limitations

Standard-GEM addresses repository-level organisation, provenance, and validation, but it does not solve all challenges associated with GEM development. First, adoption of a Git-based workflow requires familiarity with branching, commits, pull requests, and conflict resolution, which may create a barrier for some modelling groups. Second, Git is best suited to text-based files; binary formats such as .mat and .xlsx are difficult to inspect through diffs and can increase repository size. For this reason, standard-GEM encourages text-based model representations on development branches, but some binary release artefacts remain useful for compatibility with existing tools. Third, Git history is informative only when commit messages, issues, and pull requests are used carefully; poor curation practice can still produce opaque histories. Fourth, public repository hosting does not by itself guarantee long-term preservation, making archived releases through persistent repositories such as Zenodo important. Finally, standard-GEM does not assess whether a model is biologically correct. Automated validation can detect structural, formatting, annotation, and selected simulation issues, but biological validity remains dependent on evidence, expert curation, and model-specific evaluation.

### Future perspectives

At the time of publication, standard-GEM has reached version 0.9. As the field matures, it is expected that the standard will do so as well. The discussions that have already taken place in the repository indicate potential future directions, such as expanding standard-GEM to include support of multi-tissue, multi-species community metabolic models and validation improvements. Moreover, instead of becoming a static standard, standard-GEM has been set up as an “open system” to continue evolving and serving the community’s future needs. As noted by Scharm et al. (2018), “reuse of models is still impeded by a lack of trust and documentation”. Together with the standard-GEM template repository and additional resources, we will continue to support the conversion of models, as deposited in their respective publications, to the standard-GEM format. Thus, we aim to enable trustworthy and documented community curation, expansion, versioning, and transparent annotation of these models.

Future versions of standard-GEM could generalise Human2’s combination of curation assisted by large language models and repository-local automatic checks by providing optional continuous-integration workflows that evaluate proposed changes on pull requests and model performance across releases, for example by evaluation with benchmark datasets of single-gene lethality (Luo et al., 2026). Because standard-GEM provides a predictable repository structure and machine-readable validation results, these workflows could support human-supervised artificial-intelligence agents that propose evidence-linked curations as traceable pull requests and revise them in response to failed checks before expert approval, thereby closing the loop between model development and the hosted standard-GEM validation pipeline.

GEMs are no longer static outputs of individual reconstruction projects; many are continuously curated research objects that accumulate changes across years, tools, and contributors. For such models, the final deposited file captures only one state of a longer development process. This repository-centred view supports a transition from GEMs as isolated research outputs to GEMs as maintained, extensible community resources – “reusable, extensible, and published opensource” (Medley et al., 2016).

## Methods

### Development of standard-GEM

The main repository was developed by consolidating established practices for the development, versioning, and dissemination of genome-scale metabolic models into a lightweight guide. The initial specification was informed by GEM development practices at the Systems and Synthetic Biology division at Chalmers University of Technology, including experience from yeast-GEM, Human-GEM, and other community model repositories, and was further refined through public GitHub issues, internal discussions, manuscript comments, and co-author feedback. The resulting requirements and recommendations were encoded in the `.standard-GEM.md` file and implemented in a reusable template repository containing the prescribed folder structure, README template, licence, issue and pull request templates, and branch and release conventions.

### Validation pipeline

The validation pipeline starts by using the GitHub and GitLab application programming interfaces (API) to identify all standard-GEM models by finding all repositories tagged with `standard-GEM`. The pipeline then performs a validation of the repository’s content by navigating the expected file tree. After the repository has been deemed compliant, the pipeline verifies the formatting of model files and runs the model testing framework MEM-OTE (Lieven et al., 2020). Complementary efforts such as FROG analysis (Raman et al., 2024) further address numerical reproducibility of constraint-based models by standardising flux variability, reaction deletion, objective function, and gene deletion analyses into reproducible reference reports. These approaches are compatible with the modular design of the pipeline and represent possible future extensions. In addition, since previous releases of a model are all available in the same repository, tests can also be run retroactively on previous releases. Validation failures are recorded rather than used as exclusion criteria. Outputs are exported as versioned JSON files and rendered through the public website for validation reports.

## Data and code availability

Standard-GEM is available from github.com/MetabolicAtlas/standard-GEM and also at DOI 10.5281/zenodo.4447437, under the conditions of the CC BY 4.0 licence along with additional supporting material.

The output of the automated validation pipeline is available at github.com/MetabolicAtlas/standard-GEM-validation, and also at DOI 10.5281/zenodo.21039166. The documentation of the pipeline is publicly available at metabolicatlas.github.io/standard-GEM-validation/docs/.

The code to obtain the affiliation map is available at github.com/mihai-sysbio/affiliation-map.

## Author contributions

**Conceptualisation**: Mihail Anton, Dimitra Lappa, Feiran Li, Eduard J. Kerkhoven.

**Investigation**: Mihail Anton, Famke Bäuerle, Andreas Dräger, Matthias König, Feiran Li, Hongzhong Lu, Jason A. Papin, Jonathan L. Robinson, Benjamín J. Sánchez, Eduard J. Kerkhoven.

**Software**: Mihail Anton, Famke Bäuerle, Andreas Dräger, Ulf W. Liebal, Eduard J. Kerkhoven.

**Visualisation**: Mihail Anton, Andreas Dräger, Ulf W. Liebal, Eduard J. Kerkhoven.

**Writing** -original draft: Mihail Anton, Eduard J. Kerkhoven. **Writing -review and editing**: Mihail Anton, Eivind Almaas, Famke Bäuerle, Moritz E. Beber, Rui Benfeitas, Sara Benito-Vaquerizo, Lars M. Blank, Frederick Clasen, Claudia de Buck, Andreas Dräger, John M. Hancock, Vassily Hatzimanikatis, Matthias König, Dimitra Lappa, Feiran Li, Ulf W. Liebal, Hongzhong Lu, Hongwu Ma, Radhakrishnan Mahadevan, Costas D. Maranas, Adil Mardinoglu, Jens Nielsen, Juan Nogales, Marco Pagni, Jason A. Papin, Jonathan L. Robinson, Benjamín J. Sánchez, Saeed Shoaie, Vetle Simensen, Maria Suarez-Diez, Snorre Sulheim, L. Thomas Svensson, Bas Teusink, Wanwipa Vongsangnak, Ahmad A. Zeidan, Eduard J. Kerkhoven.

## ACKNOWLEDGEMENTS

We thank Iván Domenzain, Cheewin Kittikunapong, Kiran Raosaheb Patil and Hao Wang for their valuable feedback and support in developing standard-GEM.

We thank past and current members of the Systems and Synthetic Biology division at the Department of Life Sciences at Chalmers University of Technology for previous work that paved the way for standard-GEM and for valuable discussions. We acknowledge OpenAlex for providing open scholarly metadata and Natural Earth for providing public-domain vector map data used in the affiliation map.

This project was supported by the ELIXIR Systems Biology Community by way of the SYBEL Implementation Study and by the Dutch Research Council (NWO, Nederlandse Organisatie voor Wetenschappelijk Onderzoek) under grant agreement OCENW.M.21.259.

Costas D. Maranas acknowledges funding from the Center for Bioenergy Innovation supported by the U.S. Department of Energy, Office of Science, Biological and Environmental Research under Contract Number ERKP886.

During preparation of this manuscript, the authors used ChatGPT (OpenAI) to assist with language editing and editorial checks of the manuscript. All suggestions were reviewed and verified by the authors, who take full responsibility for the final manuscript.

## Supplementary Note 1: Example display of validation output

**Fig. S1.**
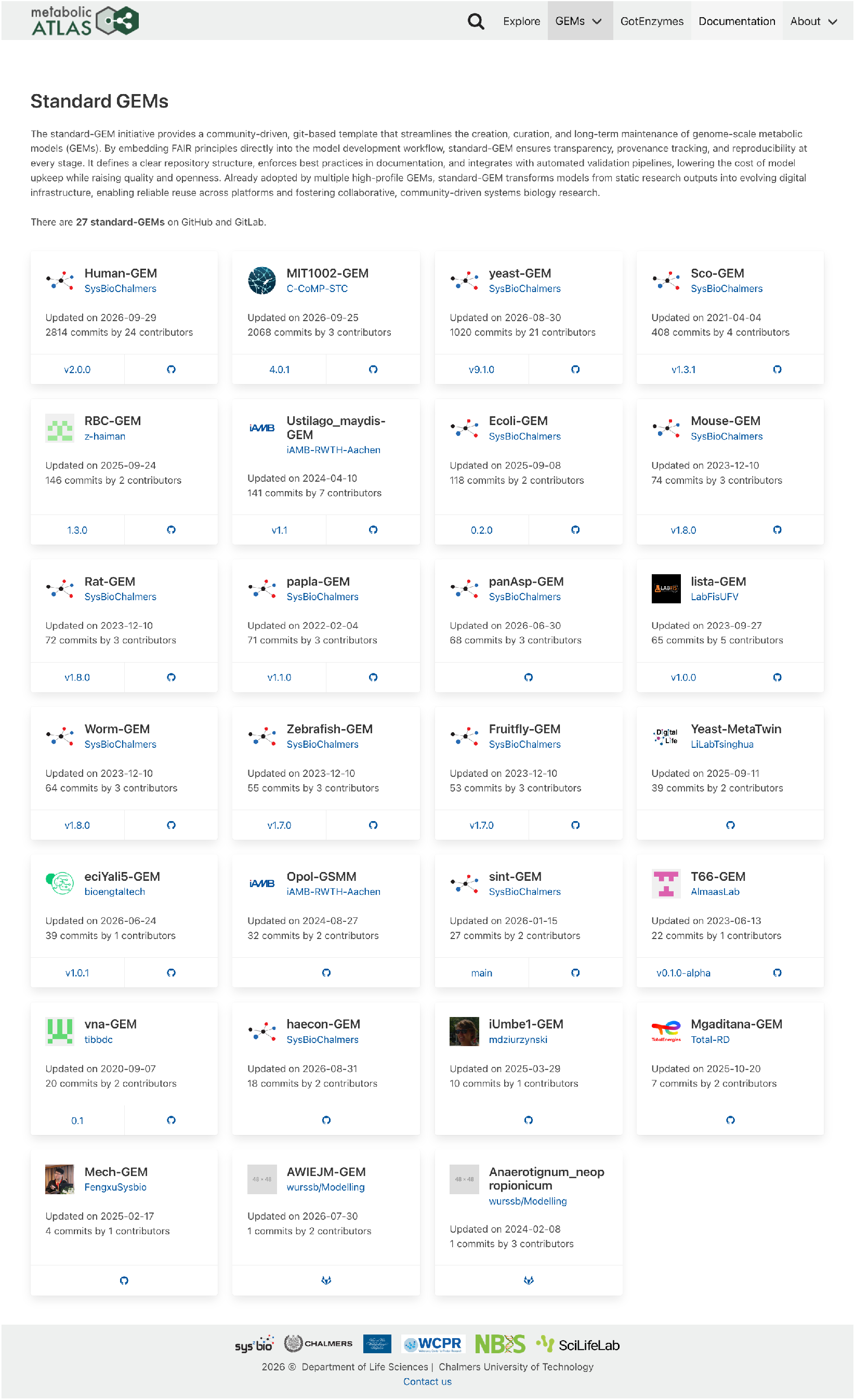
Example display of output of standard-GEM validation pipeline. This image illustrates how the validation output in JSON format can help services create pages linking to existing standard-GEMs. Screenshot retrieved 1 October 2026 from metabolicatlas.org/gems/standard-gems.

## Supplementary Note 2: Google Scholar hit-count for platforms for genome-scale metabolic modelling

**Table S1.** Google Scholar hit-count indicates that GitHub and GitLab are popular platforms for genome-scale metabolic modellers. As a rough, non-exhaustive proxy for platform visibility, we queried Google Scholar using the predefined search string `“genome scale metabolic model” [repository URL]` in June 2026. These counts should not be interpreted as exact usage statistics. Where the URL searches yielded results for both `http` and `https` protocols, both numbers are shown.

| Repository name | Repository URL | Hit count: http | Hit count: https |
| --- | --- | --- | --- |
| GitHub | github.com | 44 | 1890 |
| KEGG | www.genome.jp/kegg | 223 | 107 |
| BiGG Models | bigg.ucsd.edu | 238 | 0 |
| BioCyc | biocyc.org | 76 | 52 |
| BioModels | www.ebi.ac.uk/biomodels | 30 | 96 |
| GitLab | gitlab.com | 0 | 91 |
| ModelSEED | modelseed.org | 34 | 51 |
| VMH | www.vmh.life | 3 | 65 |
| Metabolic Atlas | metabolicatlas.org | 1 | 45 |
| MetaNetX | www.metanetx.org | 11 | 16 |
| KBase | www.kbase.us | 3 | 18 |

## Supplementary Note 3: .standard-GEM.md file

The standard-GEM template repository is centred around the .*standard-GEM*.*md* file. This file is intended to provide guidance similar to a standard operating procedure. In addition to being based on a simplified vocabulary concerning requirements, it also uses colour labels to distinguish between the three levels of requirements. For user-friendliness, checkboxes have been added to support the user through the open-sourcing process.

Specific terminology has been used in the .*standard-GEM*.*md* file. The keywords defining the standard-GEM recommendations and requirements are *must, must not, should, should not*, and *can*. These have been adapted from established requirement-language conventions, particularly RFC2119 (Bradner, 1997), and further simplified, favouring the use of *must* / *must not* over *shall* / *shall not* and avoiding the use of *may, need not* and *cannot*.

The .*standard-GEM*.*md* file is currently meant to be maintained manually. While this does include more work than using a programmatic tool which expert users might prefer, it lowers the barrier to entry for new users.

The file below was obtained from github.com/MetabolicAtlas/standard-GEM/blob/main/.standard-GEM.md.

## standard-GEM 1.0

For details about the [aims](https://github.com/MetabolicAtlas/standard-GEM/wiki/Aims,-scope-and-terminology#aims), [scope](https://github.com/MetabolicAtlas/standard-GEM/wiki/Aims,-scope-and-terminology#scope), and [use case](https://github.com/MetabolicAtlas/standard-GEM/wiki/Use-case) of this standard see the wiki pages of the standard-GEM repository.

### Terminology

The definitions used throughout this guide are copied below [from the wiki](https://github.com/MetabolicAtlas/standard-GEM/wiki/Aims,-scope-and-terminology#terminology) to facilitate understanding. Colour labels are visual aids only; the words themselves define the requirement.

Based on the ISO guidelines, tweaked for easy understanding.

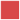 Requirements: must, must not. Mandatory for standard-GEM compliance.

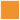 Recommendations: should, should not.

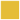 Possibility and capability: can. Optional or technically possible.

### Instructions

This document serves as a checklist for creating an open source genome-scale metabolic model (GEM) on GitHub.

□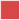All GEMs that follow the `standard-GEM` must contain this file.

The checklist records explicit confirmation by the repository maintainers. This file must be edited only with [checkmarks](https://help.github.com/en/github/managing-your-work-on-github/about-task-lists), to support automatic parsing and validation. Some of the checkmarks are pre-applied based on the contents of the `standard-GEM` template repository. GEM authors are responsible for checking that their model repository follows the guidelines entirely.

With further updates to `standard-GEM`, maintainers should replace the contents of this file with the new version, review the changes and update the checkboxes accordingly.

## Repository creation

□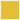Navigate to standard-GEM and click on the button `Use this template`

On GitHub, the `standard-GEM` template can be used to initiate a repository. This will copy the contents of the *main* branch into the new repository, which can be either private or public. If the repository already exists, it is sufficient to copy this file and follow the recommendations within it.

□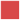Repository name

The name must be either a common name, KEGG organism, or taxonomy-derived short name, followed by the suffix `-GEM` or `-GSMM`. The `-GEM` suffix is preferred to ease pronunciation. An abbreviation can prefix the name, e.g., `ec` (enzyme constrained), `sec` (with secretory pathways), `mito` (with mitochondrion pathways), `pro` (with protein structures), or `i` (historical compatibility, see doi:10.15252/msb.20199235). Example: `ecYeast-GEM`

□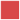The naming must be consistent.

The repository name must also be applied to the model itself, including names of model files.

□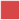Pick a repository description

The description must include the full scientific name.

Example: `The consensus GEM for Saccharomyces cerevisiae`

□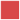Add repository topic

The topic `standard-GEM` must be added. Other topics like `genome-scale-models` and `systems-biology` can be added. Labelling the repository with this topic enables automatic discovery using the GitHub API, and automatic validation of the standard.

Repository topics are not copied when using the template and must be added manually.

□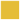Add a repository URL

The URL can be the link to the publication/preprint/website where the model is introduced, e.g., via an identifier system (doi/EuropePMC/PubMed).

## Repository workflow

□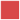Git branches

The GEM repository must have at least two branches: *main* and *develop*.

□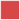Releases

Releases must use the tag format `X.X.X` where X are numbers, according to [semantic versioning principles](https://semver.org/). The last field, called “patch,” can also be used to indicate changes to the repository that do not change the GEM itself. Using a `v` before the version number (`v1.0.0`) [is discouraged](https://semver.org/#is-v123-a-semantic-version). For more information about releases, see the [documentation on GitHub](https://docs.github.com/en/github/administering-a-repository/managing-releases-in-a-repository).

□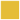*COMBINE* archive

The releases can contain a [COMBINE](https://doi.org/10.1186/s12859-014-0369-z) archive attached as a release artefact. Such an archive can be obtained in many ways, including directly from the git repository through a web service such as [CombineArchiveWeb](https://cat.bio.informatik.uni-rostock.de/#create).

□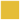Commits

Commit messages can follow the style of [conventional commits](https://www.conventionalcommits.org/).

## File tree

/signifies the root of the repository.

Git does not track empty directories. `.keep` files are placeholders that preserve the expected directory structure. They can be removed once the directory contains other tracked files.

☑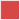`/.gitignore`

The repository must contain a `/.gitignore` file. This generic [.gitignore](https://git-scm.com/docs/gitignore) was prepared for multiple programming languages. While it does not require modification, it can be further adapted to the needs of the repository.

☑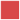`/.github`

The repository must contain a `/.github` folder, in which the contributing guidelines, code of conduct, issue templates and pull request templates must be placed. Defaults for these are provided. This folder can be named differently, to match the functionality of the Git hosting platfom.

□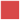`/.github/CONTRIBUTING.md`

The template provides this file, but it is empty. It must be filled in with the adequate contributing guideline instructions; a good example is

https://github.com/SysBioChalmers/yeast-GEM/blob/main/.github/CONTRIBUTING.md.

□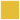`/.github/workflows`

The template can contain an optional folder for running automated workflows. The template provides a predefined workflow that runs MACAW (doi:10.1186/s13059-025-03533-6) on the YAML model file in each pull request, sharing the result as a message in said pull request, and also committing the detailed result to the repository in the `/data/testResults` folder.

□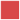`/code/README.md`

The repository must contain a `/code` folder. This folder must contain all the code used in generating the model. It must also include a `README.md` file that describes how the folder is organised.

□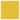`/code/test/`

The repository can contain a `/code/test` folder. All model testing code must reside in this folder.

□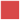`/data/README.md`

The repository must contain a `/data` folder. This folder contains the data used in generating the model. It must also include a `README.md` file that describes how the folder is organised.

□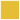`/data/testResults`

The repository can contain a folder to store the results of tests.

□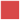`/model`

The repository must contain a `/model` folder.

This folder must contain the model files in multiple formats depending on the branch, according to the table below. As a general guideline, binary formats (`.mat`, `.xlsx`) must not exist on any other branches than *main*. The main reason is that binary files are not inspectable using text diffs, which means changes cannot be easily compared to previous versions, thus increasing the chance of errors.

Moreover, with time, the size of the repository can create difficulties, and we cannot yet recommend storing these files with Git LFS, as it introduces complexity.

For more information on the `SBtab` file format, see sbtab.net.

All model files must be named the same as the repository, and with the appropriate extension. Example: `yeast-GEM.mat`

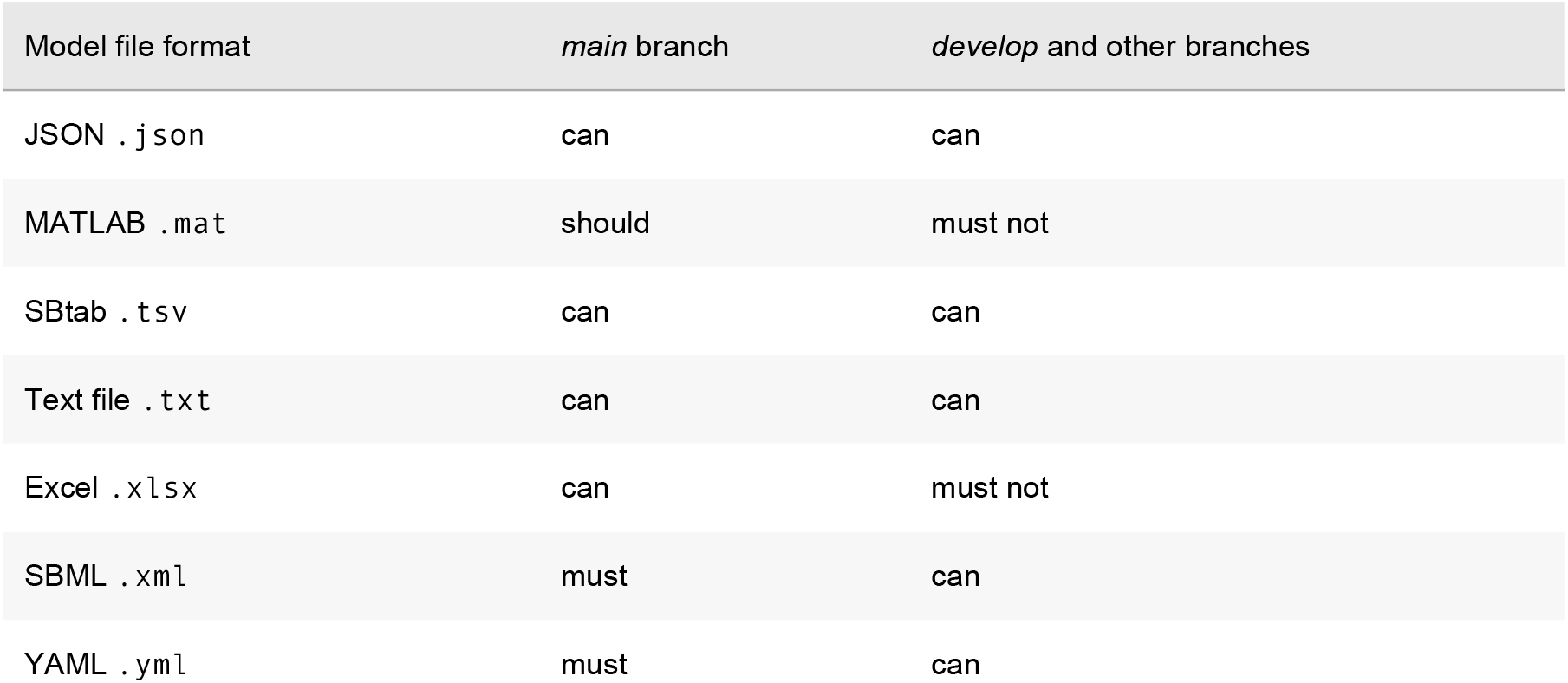

☑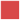`/LICENSE.md`

The repository must contain a licence file. The default licence is [CC BY 4.0](https://creativecommons.org/licenses/by/4.0/). Unless a different licence is desired, the file does not require modification.

□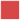`/README.md`

The repository must contain a `README.md` file. A default file is provided, and the adequate contents must be filled in.

The `/README.md` file must include a version badge. A default is provided in the file.

Additionally, the `/README.md` file should contain the [Zenodo](https://zenodo.org/) badge. After the first public release, the repository [should be archived via Zenodo](https://github.com/MetabolicAtlas/standard-GEM/wiki/FAQ#zenodo), and the corresponding badge be updated. A default is provided in the file.

The `/README.md` can contain a contact badge, e.g., [Gitter](https://gitter.io/). When setting up the Gitter chat room, the GitHub activity should be synced with Gitter in order to see the latest repository updates in the chat room. The default for this badge is provided in the file.

☑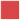`/version.txt`

The repository must contain this file, which is required for the version badge in the `/README.md`. The value refers to the GEM, not to `standard-GEM`. The value must be updated with each release.

□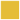Files for continuous integration testing

The repository can be set up for continuous integration testing using MEMOTE (doi:10.1038/s41587-020-0446-y) with, e.g., Jenkins (`Jenkinsfile`), and GitHub Actions (under

□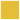.`github/workflows`).

*MEMOTE* report

The repository can contain a MEMOTE report on the *main* branch in `.html` format. The repository can be set up for continuous integration testing using memote with, e.g., Travis CI (`.travis.yml`), Jenkins (`Jenkinsfile`), and GitHub Actions (under `.github/workflows`).

